# The response to selection in Glycoside Hydrolase Family 13 structures: A comparative quantitative genetics approach

**DOI:** 10.1101/205542

**Authors:** Jose Sergio Hleap, Christian Blouin

## Abstract

The Glycoside Hydrolase Family 13 (GH13) is both evolutionary diverse and relevant to many industrial applications. Its members perform the hydrolysis of starch into smaller carbohydrates. Members of the family have been bioengineered to improve catalytic function under industrial environments. We introduce a framework to analyze the response to selection of GH13 protein structures given some phylogenetic and simulated dynamic information. We found that the TIM-barrel is not selectable since it is under purifying selection. We also show a method to rank important residues with higher inferred response to selection. These residues can be altered to effect change in properties. In this work, we define fitness as inferred thermodynamic stability. We show that under the developed framework, residues 112Y, 122K, 124D, 125W, and 126P are good candidates to increase the stability of the truncated protein 4E2O. Overall, this paper demonstrate the feasibility of a framework for the analysis of protein structures for any other fitness landscape.

## 1 Introduction

The Glycoside Hydrolase Family 13 (GH13) is a multi-reaction catalytic family of enzymes hydrolyzing α-glucoside linkages in starch. Its members catalyze hydrolysis, transglycosylation, condensation, and cyclization reactions [5]. The initial definition for this family was formulated in the early 90’s [26, 61, 34]. According to this definition, a member of this family must [61]: 1) hydrolyse or form (by transglycosylation) α-glucosidic linkages; 2) have four conserved amino-acidic regions [48]; 3) contain the catalytic triad: Asp, Glu, and Asp.; and 4) have a TIM-barrel-like fold in its structure.

Since then, the number of family members has increased [11] to include *α*-1,1-, *α*-1,2-, *α*-1,3- and *α*-1,5-glucosidic linkages [44]. Also, the number of conserved regions have been updated to 7 [29, 30]. The catalytic activity and substrate binding residues in the GH13 family members occur at the C-termini of the *β*-strands and in the loops that extend from these strands [59]. The catalytic site includes aspartate as a catalytic nucleophile, glutamate as an acid/base, and a second aspartate for stabilization of the transition state [65]. The catalytic triad plus an arginine residue are conserved in this family across all catalytic members [45, 60]. The GH13 family has many characterized enzymes with diverse functions and are summarized and clustered in the CAZy database [63]. GH13 is a highly diverse family in both function and ubiquity being found in all kingdoms of life [60]. The GH13 family has been subdivided in over 40 subfamilies [58, 33] by their sequence motif and enzyme specificity [11], but they all are related both in sequence and structure. To date, this family counts with thousands of sequences, hundreds of structures solved, and more than 30 different enzymatic specificities [11]. Many comprehensive reviews on their mechanisms, sequences, abundance, phylogeny and concept have been performed [59, 31, 43, 19, 67, 32, 10, 60].

Part of the interest in researching in this family lies in its industrial importance [22, 9], making it the target of engineering efforts to increase of thermal and alkaline stability [41, 17, 62, 21], specific activity [41, 51],, and other diverse biochemical properties that are important to the industrial context [39, 3, 17]. Many strategies have been used to engineer this family including different “rational design” approaches [12] such as B-fitter [53], proline theory [40], PoPMuSiC-2.1 [16], and sequence consensus [12]. However, to our knowledge, there is no attempt to leverage both phylogenetic and molecular dynamics signals to quantify the potential of a structure to response to selection.

Exploring how selection acts on protein structures is not a trivial problem. One approach is to assume that protein structures are shape phenotypes and that their 3D structures respond to both genetics and environmental factors, thereby falling under a quantitative genetics framework. Proteins and other shapes are highly multivariate in nature [35], and the model for their phenotype (*y*) can be expressed as [64]:

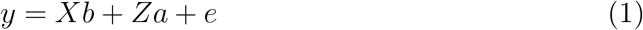

where *X* and *Z* represent design matrices for the fixed and random effects in vectors *b* and *a* respectively, and *e* is the residual component that cannot be explained by the model. Here, *y* is the phenotype of one structure and contains the x, y, and z coordinates of each homologous (with respect to the rest of the structures being analyzed) residue. For a protein structure *t* that has 100 homologous residues, the length of *y_t_* is 300. The more detailed explanation of the abstraction of the protein structure as a shape can be seen in section 2.4. With this model, the phylogenetic contribution to phenotype can be estimated. In a multivariate setting such estimation is called the *G* matrix, or genetic variance-covariance matrix, that summarizes the genetic contribution and the interaction of all traits. In the example above, *G* is a 300 by 300 matrix. Lande and Arnold [38] proposed a multivariate strategy to estimate the response to selection given *G* as:

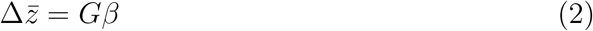

where 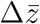 is a vector of changes in traits, and *β* is a vector of selection gradients. The latter quantity is the effect of a particular trait on the relative fitness, and therefore depends on the definition of fitness. Here we define fitness at the molecular level as the function that a particular molecule has. In enzymes, for example, this term could include the stability necessary to perform the function and the effectiveness and efficiency of the protein to do it. Then, the selection gradient can be understood as the change in fitness when the trait (in this case geometry) varies.

To apply the framework, the estimation of a G-matrix is required [36]. To deal with the fact that the number of samples is limited, this inversion of matrices require expensive computation, and an eigen decomposition of the covariance matrices is also required, the restricted maximum likelihood (REML) approach is typically employed to carry out the variance decomposition. When applied to univariate data it is more accurate than maximum likelihood methods because it better handles missing data (i.e. unknown parents, arbitrary breeding designs, etc) and can account for selection processes. However, REML has good properties only asymptotically. The reliability of the estimates is questionable when data is scarce. One way to deal with complex cases that might bias the REML estimates is to use Bayesian inference of the animal model. This approach uses Markov chain Monte Carlo simulations and is a more robust estimation than REML, with equivalent results in less complex cases [6]. This robustness assumes that the Bayesian model has enough information in the prior probability distribution. A given set of priors considerably affect the estimation of the variance components. In particular, uninformative priors, such as flat priors, can lead to biases in the estimation.

### 1.1 Lynch’s comparative quantitative genetic model: Applications to protein structures

Lynch [42] developed the *phylogenetic mixed model (PMM*). In this model, the correlation of phylogenetically heritable components is the time to the shared common ancestor (length of the path from the most recent common ancestor among two species and the root of the phylogenetic tree) in the phylogeny [24]. The PMM can be described as [42]:

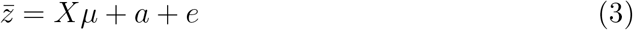

where *X* is an *np*×*p* incidence matrix, *p* being the number of traits and *n* the number of observations.

An assumption of the model is that *μ_c_* is shared among all taxa in the phylogeny. This is a sensible assumption to make when analyzing truly homologous protein structures, since the mean effect on the phenotype is shared by common ancestry. This also means that *μ_c_* + *a_ci_* can be interpreted as the heritable component of the mean phenotype for the *i*th taxon [42].

Here, the phylogenetic effects are the portion of the variation that has been inherited from ancestral species [15]. It does not only contain the genetic component, but also some environmental contributions given the shared evolutionary history of the taxa [28]. In PMM the ratio between the additive component and the total variance is the heritability (*h*^2^) in a univariate approach. Housworth et al. [28] pointed out that a univariate *h*^2^ in a PMM is actually equivalent to Freckleton et al.[20]’s and Pagel [49]’s phylogenetic correlation (λ).

Despite the robustness of the models, the REML technique, employed to estimate them, has two major drawbacks: assumption of normality of the data, and high sample size requirements. It is widely known that REML poorly estimates genetic correlation when overparameterized (multi-trait inference), when the sample size is small (Martins, personal communication), and when the normality assumption is violated [24]. These violations can be handled in a Bayesian framework using Markov Chain Monte Carlo techniques. In such techniques, the higher complexity of the joint probability calculation needed for the likelihood estimation can be broken down in lower dimensional conditionals. From those conditionals the MCMC sampling can be performed and marginal distributions can be extracted [24]. A discussion of the use of Bayesian MCMC techniques is beyond the scope of this work. We refer the interested reader to Sorensen and Gianola [57] for a good description of likelihood and Bayesian methods in quantitative genetics.

Despite its strengths, the Bayesian framework also has weaknesses. The most important one is that it requires proper and informative priors. Uninformative priors lead to biases with high variation in results. The sensitivity to the choice of prior distribution should always be assessed [37]. Given that in evolutionary biology datasets the amount of knowledge on the estimator is scarce, well informed priors are normally not available and by informing priors with partial information, the estimation can become ill-conditioned.

To explore the feasibility of a comparative quantitative genetics (CQG) framework in protein structures, we simulated a dataset with variable numbers of traits and observations. We show that the current implementations of the CQG framework are not feasibly applied to the dimensionality required for protein structures. We devised a method that functions as a proxy for the CQG framework and show that it is feasible and accurate. By applying this framework using the energy of unfolding (Δ*G*°) as fitness function to the GH13 family, we are able to show how purifying selection have fixed the geometry of the TIM-barrel. We also demonstrate how by changing the fitness function, the response to selection propensity changes accordingly. Finally, a proxy for the amount of dynamic deformation happening in the protein given a vector of selection is explored. Overall, we present here a starting framework to explore protein structure evolution and design.

## 2 Methods

### 2.1 GH13 dataset

Given that molecular dynamic simulations are very time consuming, we used a subset of the proteins classified as Glycoside Hydrolases Family 13 (GH13). We randomly selected 35 protein structures from a possible set of 386, but one failed during the MD simulation. A final set of 34 protein structures (Table A1 in supplementary methods) was used in further analyses.

### 2.2 Molecular Dynamics (MD) simulations

Each of the 34 protein structures was simulated in solution using the software `GROMACS 4` [27]. The force field modes used for the simulations were GROMOS96 for the protein, and the SPCE for the water molecules. Data were collected every two picoseconds for at least 40 nanoseconds, discarding the first 10 nanoseconds of simulation to achieve stability. This process was performed using a workstation with 24 CPU cores and an NVIDIA TESLA^TM^ GPU.

The analysis of these simulations will provide information on the flexibility (or within protein variance) of the protein, as opposed to the analysis across homologs that will provide phylogenetic information (or between structures variance). By 40 ns all proteins analyzed have achieved equilibrium and therefore most of the intrinsic variance has been captured.

### 2.3 Aligning the structures and MD simulations

The alignment of homologous proteins was performed using `MATT` software [47]. To align the snapshots from MD simulations a General Procrustes Superimposition (GPS) was performed using the `R` package `shapes` [18].

### 2.4 Abstracting protein structures as shapes

On a set of aligned protein structures, the abstraction is performed in a similar way to that in Adams and Naylor [1]. However, they do not fully describe the abstraction.

Here we assign a landmark to the centroids of residues defined by:

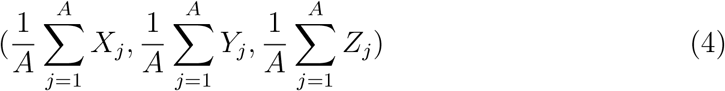

where *A* will be the number of heavy atoms (C, O, N) that constitute the side chain of a residue including the alpha carbon (*C*_*α*_). This procedure takes into account only the homologous residues. It captures the variance of both the backbone and the side chain. In the case of glycine, the centroid is the *C*_*α*_.

Once the structure is abstracted as a shape, the resulting *n* (number of observations) by *l* (number of coordinates of homologous residues) matrix is referred to as the phenotypic matrix (*P*). For example, let us assume that we have a protein structure composed of 150 residues. Let’s imagine that 100 different taxa share an ortholog of this protein. After aligning the protein structures let’s assume that 100 residues are homologous across all 100 taxa. The resulting phenotypic matrix (*P*) will be composed of 100 rows of observations (*n*) and 300 coordinates. These dimensions correspond to the x, y, and z axis of each of the 100 homologous residues. To estimate the variation of this phenotype, the phenotypic variance *V_P_* can be estimated by computing the variance-covariance matrix of *P* as *V_P_* = *var*(*P*), or *G* in a multivaraite scenario.

### 2.5 Pooled-within group covariance matrix estimation

After the MD simulations up to 500 samples per simulation were obtained. The estimation of the pooled-within covariance matrix was performed as follows:

1. Align every model within each MD simulation using General Procurstes Superimposition (GPS): Remove extra rotations and translations that could occur during MD simulation.
2. Select an ambassador structure that is closest to the mean structure (the geometrical mean of the dataset).
3. Align all ambassadors using MATT flexible structure aligner to identify homologous sites: Multiple structure alignment to identify structural homology.
4. Extract the centroid of fully homologous sites: Identify shared information among all structures.
5. Concatenate the centroids’ three dimensions for all trajectories
6. Perform a GPS on the entire set of shapes to bring all pre-aligned structures into the same reference plane.
7. Compute the pooled-within covariance matrix (*W*) by first computing the deviation from the mean in each class/group (individual homologs in our case) as:

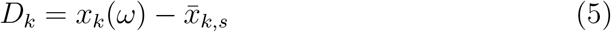

then computing the sum over the classes of the products of *D_k_* as:

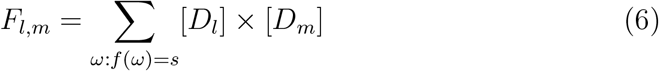

Finally, compute the pooled-within covariance matrix:

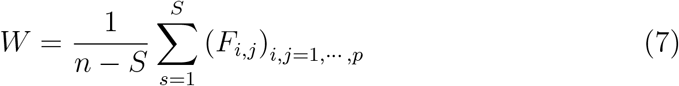

where *S* is a the number of categorical variables describing the groups or species, *ω* is an instance where *f* (*ω*) corresponds to the class value of the instance, and 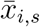 is the mean of the variable *i* for individuals belonging to *s*. Finally, *n* is the sample size.

Here, *W* contains the covariance matrix of the within-homolog (i.e. Molecular dynamic data). To estimate the evolutionary component of *P*, the between structures/species covariance matrix (*B*) has to be taken into account. *B* will be simply the difference between the *V_P_* and *W*.

### 2.6 Estimating 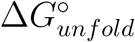 as proxy for fitness

The 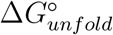 on each model for each protein was estimated using the command line version of `FoldX` [56]. It is important to notice that the computed 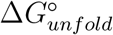 is not comparable in proteins of different size, therefore we computed the average 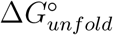 per residue as:

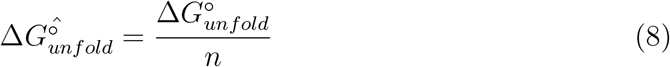

*n* being the number of residues. With this 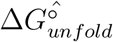 as proxy for fitness we can try to explore the fitness surface. To do this, we used the first two principal components (PC) of a PC analysis of the shapes as X and Y axes; 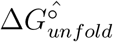 in the Z axis (Supplementary figure B3).

### 2.7 Propensity to respond to selection

Arnold [4] showed that, despite high additive variances, *G* might not be aligned with the fitness surface. This implies that even though *β*_λ_ can be non-zero, the response to selection might send the phenotype in a different direction than the fitness surface. Blows and Walsh [8] and Hansen and Houle [25] developed an approach to measure the angle between *β* and the predicted response to selection from the multivariate breeders equation, 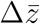 as:

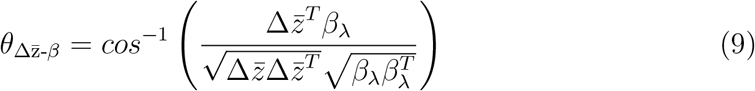

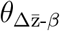 would be zero when there is no genetic constraint, whereas an angle of 90° would represent an absolute constraint [66].

## 3 Results and Discussion

In supplementary materials A and B we have shown that the traditional PMM models and their Bayesian counterparts are not feasible when the number of traits and observations are in the order of those obtained in protein science when MD simulations are taken into account. Here, we applied a simple method to overcome this over-parameterization.

### 3.1 Overcoming over-parameterization: Approaching the G-matrix by means of the P-matrix

Given the previous results, the estimation of the G matrix within the Lynch’s PMM is not feasible. This is not a new observation since in comparative evolutionary biology it is widely known that accurate measures of *G* are difficult or impossible to obtain [46]. This pattern is even more evident when dimensionality is high. On average, protein structures are composed of over 200 residues in a three-dimensional system, which means over 600 variables. Also, the sample size at the species level is typically small. Because of these reasons, a full and stable estimation of the *G*-matrix is not possible. However, an increased number of samples can be achieved by means of molecular dynamic simulations. This increases *n* considerably depending on the length of the simulation. We have shown the infeasibility of the GLMM to deal with the dimensionality and very large sample size. However, it has been shown that phenotypic (*V_P_*) covariance matrices can be estimated with more confidence with large sample sizes [13]. It is also shown that in some cases, *V_P_* can be used as surrogate for *G* when the two are proportional [46, 55]. To test this, we performed a shape simulation explained in supplementary section A.1. The simulation was performed with 500 replicates as molecular dynamics snapshots, 100 taxa, and the traits were varied from 2 to 1024 in a geometric series increase. Since the within-homolog matrix structure is known, a pooled-within covariance matrix (*W*) was computed as exposed in the section 2.5.

Table 1 shows the feasibility and accuracy of the pooled-within species covariance estimation method. Here the Cheverud’s Random Skewer (RS) test [13, 14] implemented in the `R` package `phytools` [54] were used to test the accuracy. A discussion of the appropriateness of the usage of this metric can be found in Supplementary Materials A3 and references therein.

**Table 1:**
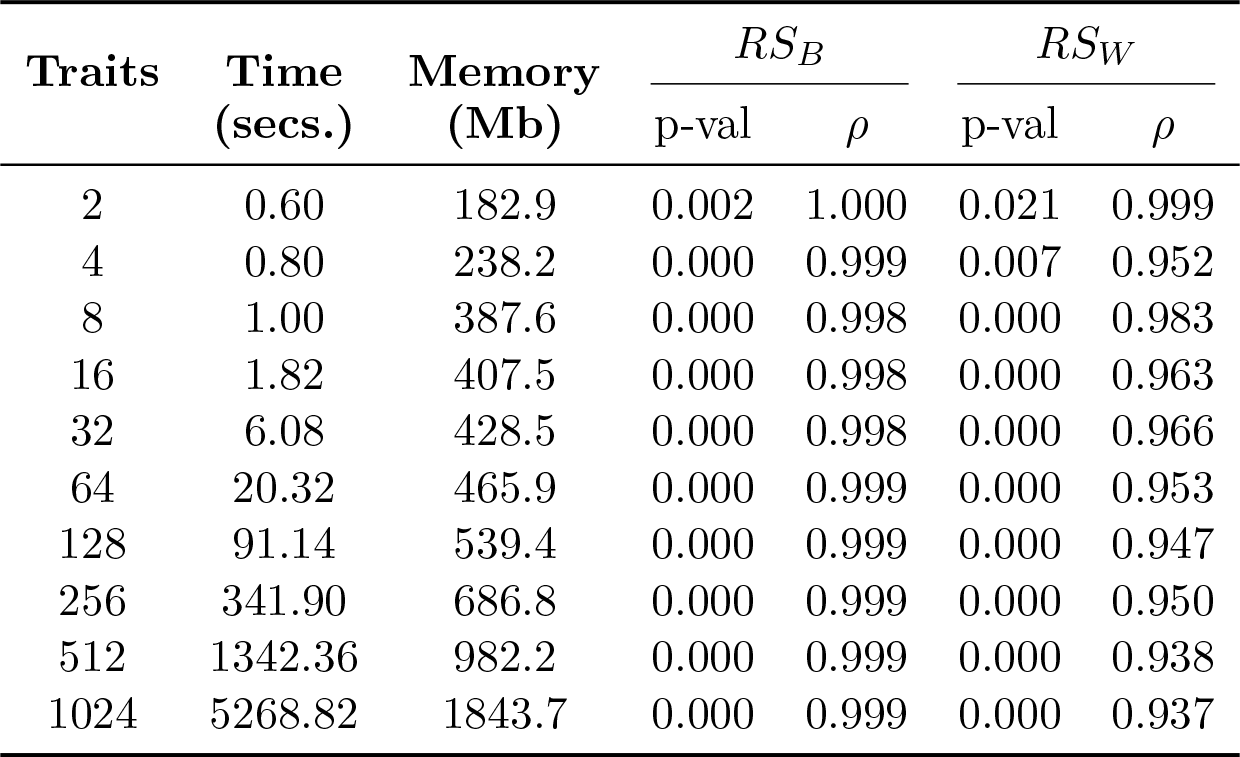
Accuracy and feasibility of the pooled-within covariance estimation. Memory (Mb), time (sec) and accuracy (random skewer correlation) of the pooled-within covariance estimation approach. *RS_B_* corresponds to the random skewer test for the phylogenetic covariance and *RS_W_* to the dynamic component.

Even with highly multivariate data (1024 traits), the memory requirement is manageable (less than 2 Gb), the evaluation is completed in under an hour, and the accuracy of the estimation is high. The estimated *G* matrix is almost identical to the simulated one is most of the runs, and the estimated *MD* have over 0.97 correlated responses to random vectors than the actual *MD*. This is a surprising result since this method cannot completely separate the error terms from the genetic and the dynamic components. However, the split of the error term between the two other components can make it negligible. Moreover, it seems that error does not significantly affect the structure of *G* and *MD*, allowing them to behave almost identically in comparison to the simulated counterparts. Given these results, and the fact that the application to real datasets can only be made with this approach, it is reasonable to keep using the described method from this point forward. However, the biological and evolutionary meaning of this approach is less clear than in the other methods since there is no explicit use of a phylogeny.

#### 3.1.1 Meaning of the pooled within-structure covariance matrix

*V_P_*-matrices can be used as surrogates of G-matrices in cases were they are proportional or sufficiently similar [50]. Proa et al. [50] showed that this assumption can be relaxed if the correlation between *G* and *V_P_* ≥ 0.6. In protein structures, we can assume that given the strong selective pressures and long divergence times, the relationship between *V_P_* and *G* is standardized. Assuming that this is true in protein structures, the estimated pooled variance-covariance (V/CV) matrices in real datasets might have a specific biological meaning. This was described in Haber [23] for morphological integration in mammals. Following Haber’s [23] logic, the within-structure/species (i.e. thermodynamic V/CV) matrix refers to integration of residues in a thermodynamic and functional manner. It also contains information about environmental factors affecting the physical-chemistry of the structure. Haber [23] includes a genetic component for his estimation of the within population variation, since populations follow a filial design. Our data, on the other hand, have a controlled amount of genetic component given that the sampling is done in a time series instead of a static population. Our approach would be more related to an estimation of within repeated measures design.

The among-structure/species (i.e additive or evolutionary V/CV) matrix refers to the concerted evolution of traits given integration and selection [23].

### 3.2 Response to selection in the GH13 family

As defined in equation 2, the response to selection of a phenotype depends on the within-species change in mean due to selection, the correlation between different traits, and the amount of heritable component of the shape. The first component can be referred to as 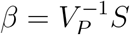, and also known as the vector of selection gradients [52] or directional selection gradient. The second and third elements are summarized in the *G* matrix. As expressed in equation 2, this covariance matrix represents the genetic component of the variation in the diagonal, and the correlated response of every trait to each other in the off-diagonal.

Another extension from equation 2 is to compute the long-term selection gradient assuming that *G* is more or less constant over long periods of time:

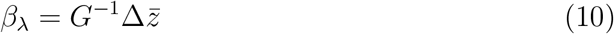

Here 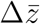 would be proportional to the differences in mean between two diverging populations.

It is important to stress the relationship between these concepts and fitness. Given that fitness (*w*) is directly related to selection, its mathematical relationship can be expressed as 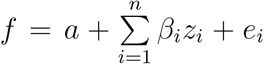 [8], and so it behaves as the weight of a multiple regression of *f* on the vector of phenotypes *z*.

In proteins, the definition of fitness is not trivial, and can vary depending on the hypothesis being tested. If the analysis is done comparatively (i.e. across different protein structures from different sources), a fitness analysis including exclusively structural measures, such as Gibbs free energy (Δ*G*), can be misleading. The fitness surface that can arise from this data would only represent departures from every individual native state. Nevertheless, Δ*G* and the energy of unfolding (Δ*G*°), are important measures to determine the stability of the protein which is important for the fitness of a protein structure. The stability of the structure allows it to perform a function and is therefore under selection because it is necessary for the particular biochemical function [7]. We are aware that there is a limitation to the protein structure stability role in fitness. To improve this fitness landscape, *f* can be defined by Δ*G*° coupled with a functional measure. In proteins, function is the main selective trait; therefore, including a term accounting for this would create a more realistic fitness surface. In enzymes this can be achieved by using the *K_cat_*/*K_M_* for each of the enzymes for a common substrate. The fitness function (*F*) can be expressed as:

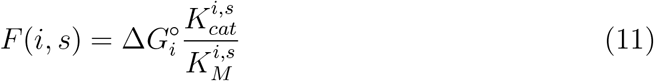

where 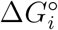 is the free energy of unfolding of the structure *i*, 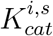 is the turnover number for structure *i* in substrate *s*, and 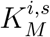 is the Michaelis constant of protein *i* working on substrate *s*.

In the case of the α-amylase family (GH13), one might try to apply the framework developed in previous sections and try to estimate the response to selection of a subset of them. However, equation 11 cannot be applied since the information of the relative efficiency given a common substrate is not consistently available across all proteins in the dataset. For this reason we are going to work exclusively with 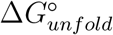, keeping in mind two caveats, 1) that 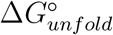 only represents structural stability and 2) that it has been shown that Δ*G_equilibrium_* or 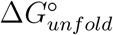 are not optimized for during evolution [2].

#### 3.2.1 Estimating dynamic and genetic variance-covariance matrices in the *α*-Amylase dataset

The structure depicted with the higher fitness was the model 1 of structure 2TAA (Supplemental figure B3), from *Aspergillus oryzae* assuming 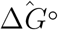 as fitness. The model 1 of structure 2TAA can be assumed to be the result of the goal of selection. The realized response to selection 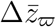 can be defined as *μ*_⊕_ - *μ*_0_, where *μ*_⊕_ is the target or after-selection mean structure and *μ*_0_ is the starting or before-selection structure. To estimate 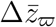 it is essential to have the fitness defined based on the questions to be asked, given that the interpretation of the realized response to selection depends on it.

In an engineering perspective, let’s assume that *μ*_⊕_ is the mean of a population of structures with the desired stability. On the other hand, *μ*_0_ is the mean of a population of structures created by a desired vector. One might ask the question of how does *μ*_0_ have to change towards the stability of *μ*_⊕_. This can be achieved by computing *β*_λ_ (equation 10), and replacing 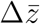 by 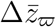. In the particular case of the GH13 dataset, let’s assume that the model 1 of the structure 2TAA is the desired phenotype (with the higher fitness in supplementary figure B3), and the model 643 of the structure 4E2O from *Geobacillus thermoleovorans* CCB_US3_UF5 (with the lower fitness in supplementary figure B3) corresponds to the source phenotype. *β*_λ_ would have a length corresponding to the dimensions of the shape. In the GH13 case 297 homologous residues were identified, which means that these shapes have a dimensionality of 891 traits. This dimension-per-dimension output is important since it reflects the amount of pressure in each dimension per each residue. However, it makes the visualization more difficult. For the sake of visualization simplicity, Figure 1 shows the absolute value of the sum of *β*_λ_ per residue, standardized from 0 to 1.

Figure 1a shows the selection gradient using the estimated *G*. Not surprisingly, the selection gradient for the TIM-barrel is low. This means that there is not much directional selection on this sub-structure. However, it is somewhat surprising that there is not any purifying selection either. This can be explained by the fixation of the trait in the evolution. Since the TIM-barrel is a widespread sub-structure that has been strongly selected during evolution, it might have reached a point of fixation of its geometry. Therefore, the *G* matrix shows little covariation among these residues since the geometric variability is also low. It is important to stress here that the phenotype measured is the geometry of the structure more than that of the sequence. Therefore, despite some variation that may have occurred at the sequence level, it might not have meaningfully affected the positional information.

**Figure 1:**
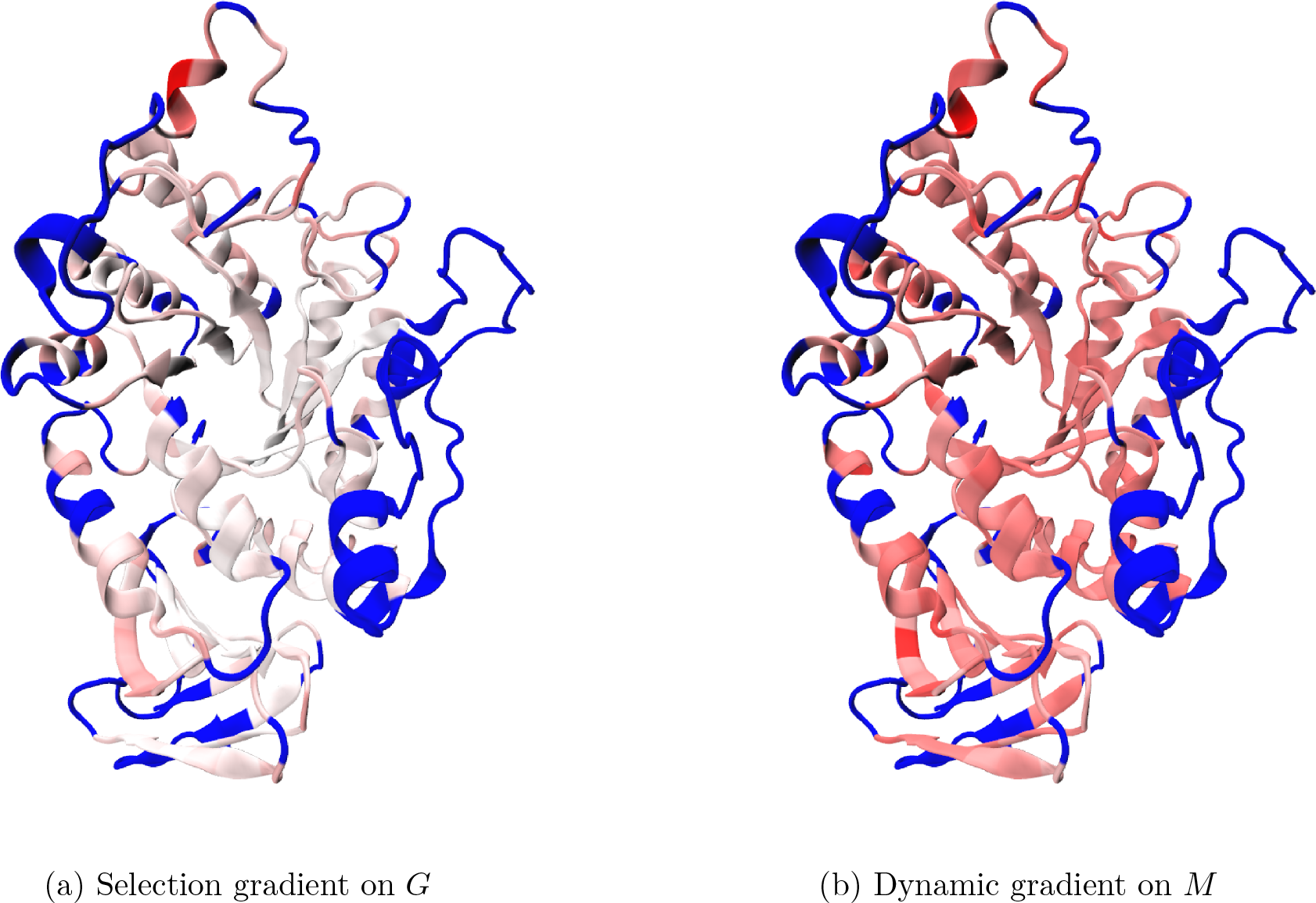
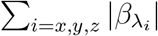 rendered in the source structure 4E2O. White represents the lowest magnitude (0), while red the highest (1). Blue depicts the non-homologous residues.

However, one must be cautious with the approach employed in Figure 1 since the signs are missed, thereby ignoring the direction of selection and the correlated response to selection. Nevertheless, this approach allows for a coarse-grained visual exploration of 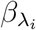. Individual instances identified by this method should be analysed afterwards in each dimension. Table 2 shows the actual values of *β*_λ_ for the top 5 positive values (directional selection) and top 5 negative values (purifying selection).

**Table 2:**
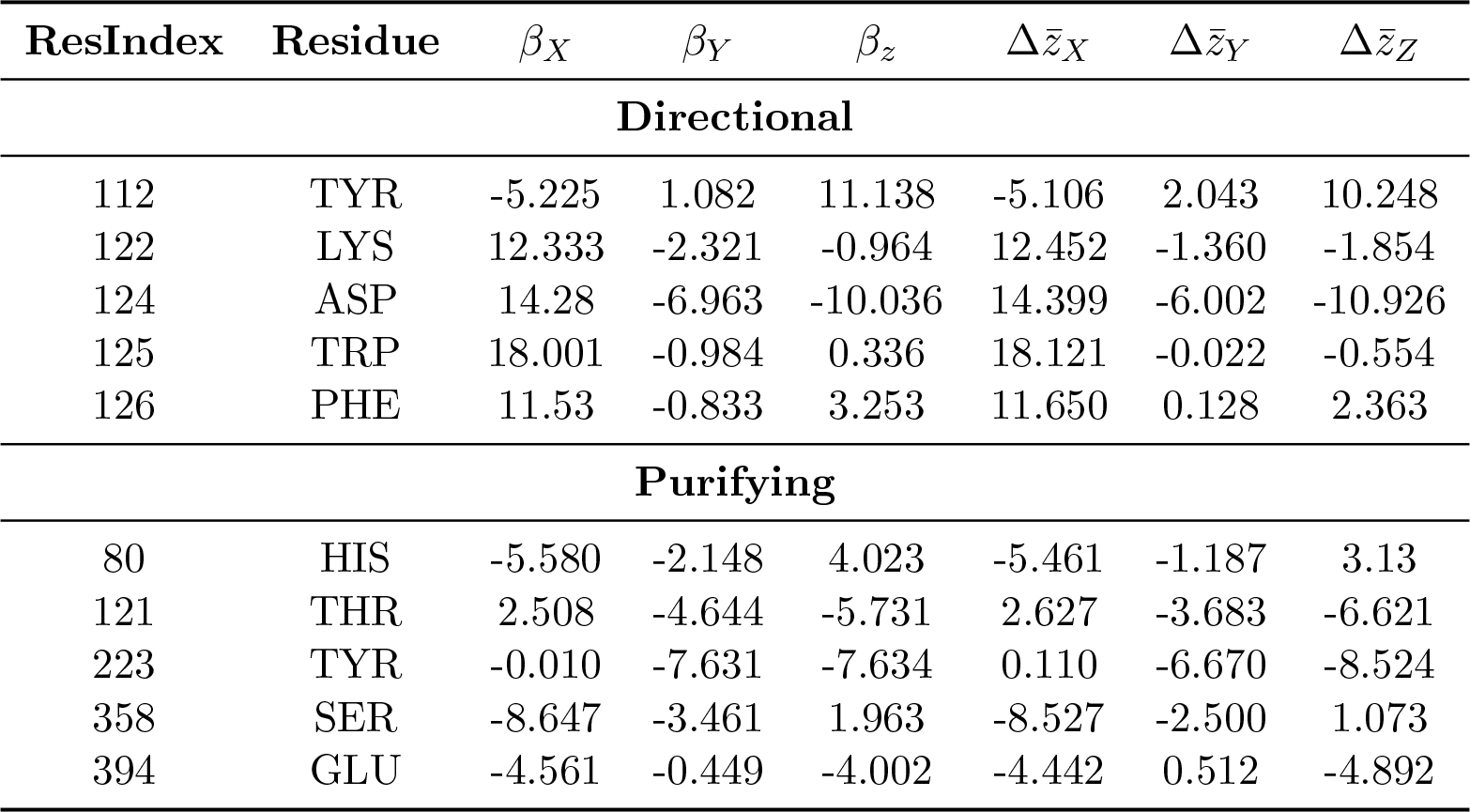
Selection gradient in the top 5 residues. Top panel shows the residues were at least one of its coordinates is under directional selection and the sum of their absolute values is the highest. Bottom panel contains the information of residues where at least one of its coordinates is under purifying selection, and the sum of the raw values are the lowest.

Figure 1b and Table 3 show the mean difference between target and source when effects of correlated dynamic differentials are removed. Given that effectively *G* acts as a rotation matrix in equation 10 to remove the selection differentials, one may posit that the same can be achieved with the dynamic (*M*) matrix. This concept is more difficult to interpret than the actual response to selection. Once *G* is replaced by *M* in equation 10, we might call it *dynamic gradient* to differentiate it from the selection gradient already explained. In this case, if the gradient is zero for a given trait, this can be interpreted that the dynamic component of the phenotype does not contribute significantly to the difference in shape for that particular trait. In the case of non-zero gradients, these can be interpreted as contributions of the dynamics to the differential, either towards the target (positive gradient) or away from the target (negative gradient).

**Table 3:** Dynamics gradient in the top 5 residues. Top panel shows the residues where at least one of its coordinates is under positive gradient. Bottom panel contains the information of residues where at least one of its coordinates is under a negative gradient.

| ResIndex | Residue | $\beta_X$ | $\beta_Y$ | $\beta_z$ | $\Delta\bar{z}_X$ | $\Delta\bar{z}_Y$ | $\Delta\bar{z}_Z$ |
| --- | --- | --- | --- | --- | --- | --- | --- |
| **Directional** |  |  |  |  |  |  |  |
| 117 | LEU | 13.028 | 37.149 | 11.848 | 2.130 | 3.521 | 4.437 |
| 125 | TRP | 29.019 | 33.605 | 6.857 | 18.121 | -0.022 | -0.554 |
| 126 | PHE | 22.548 | 33.755 | 9.774 | 11.650 | 0.128 | 2.363 |
| 262 | LYS | 12.972 | 38.081 | 11.412 | 2.073 | 4.454 | 4.001 |
| 367 | LEU | 13.590 | 34.561 | 15.609 | 2.692 | 0.933 | 8.197 |
| **Purifying** |  |  |  |  |  |  |  |
| 124 | ASP | 25.297 | 27.625 | -3.515 | 14.399 | -6.002 | -10.926 |
| 223 | TYR | 11.008 | 26.958 | -1.113 | 0.110 | -6.670 | -8.524 |

In the GH13 subset, most dynamic gradients were positive having only two residues that had one coordinate under a negative gradient (Table 3). This can also be inferred by Figure 1b. The values of the dynamic gradient are high but sensible given the definition of fitness. Since we defined fitness as the energy of unfolding (Δ*G*°), most of the information used to select the target and source structures comes from stability, and therefore thermodynamic information. The results depicted in Table 3 and Figure 1b suggest that most of the variation that explains the difference in phenotype between the structure 4E2O and 2TAA, is contained within the molecular dynamic component rather than the approximation to the phylogenetic component.

##### Orientation of *G*

The GH13 *θ* was 1.4 degrees, which means that the direction of optimal response is 1.4 degrees away from the total genetic variation of 99% explained by the projection. According to this, the *Geobacillus thermoleovorans* structure is susceptible to the selection in the actual direction of the fitness landscape towards the structure of *Aspergillus oryzae* to achieve maximum stability. The extent of such change is given by 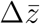, which means that the centroid position of the residue *i* should be displaced by 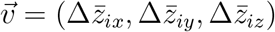.

In the case of the dynamics, the same approach can be taken. Here, *θ_M_* was 1.5 degrees which means that the optimal dynamic response is 1.5 degrees away from the optimal response. This can be interpreted in a similar way than that of the regular *θ*. However, manipulating the structure along the dynamics gradient is not feasible.

The GH13 dataset 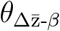 was 0.3. This means that the genetic constraints on 4E2O are not affecting the direction of selection. This posits the possibility that a strong directional selection will drive the source structure towards the target. The same pattern happens when this approach is applied to *M*. 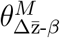 is 1.46 degrees, which is almost identical to *θ_M_*. Thus, there are almost no within-variation or dynamic constraints to the vector of response given the dynamic gradient.

## Concluding remarks

We have introduced the application of the approximation of comparative quantitative genetics framework, by means of a pooled-within group covariance matrix in a subset of the GH13 proteins, and demonstrated this application is feasible and provides sensible results, given the definition of fitness. This definition is essential in the interpretation of the results since it is the interpretation that gives polarity to 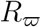. Therefore, all conclusions about the response to selection and the selection gradient itself must be analyzed under this light.

The usage of *M* in the determination of the dynamic gradient could be controversial. This is due to the fact that, in the partition of the phenotypic variance, *M* is expected to be the environmental variance plus an error term. However, since the source data for the estimation of *G* and *M* come from repeated measures by MD, *M* contains information about the thermodynamics and folding stability of the protein. It is therefore also contributing to selection.

It is important to stress the fact that this is an approximation to the true *G* and true *M*, since we have shown in previous sections that these cannot be estimated given the dimensionality of the phenotype. However, we have shown that the pooled-within group approach gives consistent results.

We have also shown that, in a stability perspective, the TIM-barrel show a small phylogenetic/genetic component to the selection gradient when a less stable structure (4E2O) is analyzed with respect to a more stable one (2TAA). In an engineering perspective, this means that most of the changes in shape come from the dynamics. Nevertheless, the small 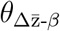 show that most of the changes applied to 4E2O would directly result in increasing the stability towards the one expressed by 2TAA. 4E2O is a truncated protein, and therefore some loss of stability is expected. It seems that residues 112Y, 122K, 124D, 125W, and 126P, are good candidates to increase the stability of the molecule given their 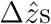. In these cases, the goal will be to shift the position of their centroids by the resulting vector of the three dimensions.

### A Material and Methods

This section contain all the information on the structures used. It also have all the simulation and test methods performed to show the infeasibility of the traditional and bayesian PMM in protein structures. This supplementary material can be found in here

### B Supplementary results

All of the simulation and test results showing the infeasibility of the traditional PMM in protein structures can be found in here

## Acknowledgments

The authors thank the members of the Blouin Lab for helpful comments and critical review of this manuscript. We also thank Jitka M. Krejci for the editorial and language revision of the manuscript. This study was funded by NSERC through the grant No. 120504858. This work was partially supported by the Departamento Administrativo de Ciencia y Tecnología - Colciencias (Colombia) through the CALDAS scholarship.

